# Chromatin remodeller *Chd7* is developmentally regulated in the neural crest by tissue-specific transcription factors

**DOI:** 10.1101/2022.11.03.515045

**Authors:** Ruth Williams, Güneş Taylor, Irving T C Ling, Ivan Candido-Ferreira, Sarah Mayes, Yavor K Bozhilov, Richard C V Tyser, Shankar Srinivas, Jim R Hughes, Tatjana Sauka-Spengler

**Affiliations:** University of Oxford, MRC Weatherall Institute of Molecular Medicine, Radcliffe Department of Medicine, Oxford, UK; Stowers Institute for Medical Research, Kansas City, MO, USA; University of Oxford, Department of Paediatric Surgery, Children’s Hospital Oxford, Oxford, UK; University of Oxford, MRC Centre for Computational Biology, MRC Weatherall Institute of Molecular Medicine, Oxford, UK; University of Oxford, MRC Molecular Haematology Unit, MRC Weatherall Institute of Molecular Medicine, Radcliffe Department of Medicine, Oxford, UK; University of Oxford, Department of Physiology, Anatomy and Genetics, South Parks Road, Oxford, UK; Francis Crick Institute, Stem Cell Biology and Development Genetics Laboratory, London, UK; Janssen Research and Development, 1400 McKean Rd, Spring House, PA, USA; Wellcome-MRC Cambridge Stem Cell Institute, University of Cambridge, Cambridge, UK

**Keywords:** Chd7, neural crest, CHARGE syndrome, enhancer, chicken

## Abstract

Neurocristopathies such as CHARGE syndrome result from aberrant neural crest development. A large proportion of CHARGE cases are attributed to mutations in the gene encoding *CHD7*, chromodomain helicase DNA binding protein 7, which remodels chromatin. While the role for CHD7 in neural crest development is well documented, it remains elusive how this seemingly ubiquitous factor is specifically upregulated in neural crest cells. Here, we use epigenomic profiling of chick neural crest to identify a cohort of enhancers regulating *Chd7* expression in neural crest cells and other tissues. We functionally validate upstream transcription factor binding at candidate enhancers, revealing novel epistatic relationships between neural crest master regulators and *Chd7*. To our knowledge, this is the first report of tissue-specific regulation of a chromatin remodeller. Furthermore, we find conserved enhancer features in human embryonic epigenomic data and validate the activity of the human equivalent *CHD7* enhancers in the chick embryo. Collectively our findings embed *Chd7* in the neural crest gene regulatory network and offer potentially clinically relevant elements for interpreting CHARGE syndrome cases without causative allocation.

## Introduction

The neural crest is a transient and migratory embryonic progenitor population that contributes derivatives to a remarkable range of neural and mesenchymal tissues in the vertebrate body. Neural neural crest derivatives include craniofacial ganglia, and both neurons and glia of the peripheral and enteric nervous systems. Mesenchymal neural crest derivatives include craniofacial cartilage and bone, as well as smooth muscle of facial blood vessels, striated muscle forming the cardiac outflow tract and septum, and the majority of body’s pigment cells. Consequently, errors in neural crest patterning, migration and differentiation result in a wide range of congenital birth anomalies collectively termed neurocristopathies. Neurocristopathies account for nearly one third of all birth defects (1). These include CHARGE syndrome, which affects the eye, heart and facial structures(2, 3), Hirschsprung’s disease, characterised by the loss of neural crest-derived enteric ganglia (4, 5), Waardenburg syndrome, characterised by deafness, pigmentation and craniofacial defects (6), and Treacher Collins Syndrome presenting with craniofacial defects (7, 8). Neurocristopathies can be caused by mutations in master neural crest regulators, for example, *SOX10* in Hirschsprung’s disease and Waardenburg syndrome (9–13) or *PAX3* in Waardenburg syndrome (14–16). However, mutations in genes encoding general cellular machinery can also result in neurocristopathies as demonstrated in Treacher Collins Syndrome caused by mutations in RNA polymerase I (8, 17, 18) and CHARGE syndrome where heterozygous mutations in the chromatin remodeller *CHD7* (chromodomain helicase DNA binding protein 7) are detected (19, 20).

CHARGE syndrome (OMIM 214800) patients present with ocular Coloboma, Heart malformations, chonanal Atresia, Retardation of growth, Genital hypoplasia and Ear abnormalities. Over 500 different pathogenic mutations in the *CHD7* coding region have been described, accounting for 32% to 41% of CHARGE syndrome cases (21). Mutations have been reported throughout the *CHD7* gene body, indicating premature termination of the protein is significantly detrimental to CHD7 function (20–25). CHD7 influences gene regulation (21, 26) by catalysing nucleosome repositioning in an ATP-dependant manner (27). In keeping with this regulatory role, Chd7 associates with distal regulatory sites carrying H3K4me1 chromatin modifications characteristic of poised enhancers, in mouse embryonic stem cells (28, 29).

While CHD7 is part of the omnipresent chromatin remodelling machinery and is broadly expressed in a multitude of embryonic tissues, the clinical features of CHARGE syndrome clearly suggest CHD7 has tissue and developmental stage-specific roles (3). Previous studies have shown that CHD7 function is essential for proper neural crest development and migration. In human neural crest-like cells, CHD7 occupies distal regulatory elements for neural crest transcription factors *SOX9* and *TWIST1* (29). In mice, *Chd7* heterozygotes present with CHARGE syndrome-like features (28, 30, 31) and trunk neural crest cells require Chd7 to maintain their multipotency (32, 33). In Xenopus, *Chd7* mutant embryos have reduced *Sox9, Twist1* and *Snai2* expression and display CHARGE syndrome like features (29). Recent work in the chick neural crest, employing weighted gene co-expression network analysis (WGCNA) analysis (34), showed that *Chd7* expression strongly correlated with expression of neural crest regulators (*Sox5, Sox9, Zeb2* and *NeuroD4*), other chromatin remodellers (*Kdm1B, Kdm2A, Kdm3B, Kdm7A*) and Semaphorins (*Sema3A, Sema3E, Sema4D, Sema6D*) which have previously been shown to be regulated by Chd7 in mice (35–37). Collectively, these findings provide strong argument for positioning *Chd7* within the neural crest gene regulatory network. However, the identity of tissue-specific enhancers and upstream neural crest factors mediating tissue specific enrichment of *Chd7* in the neural crest have not been investigated.

Here we identify super enhancer-like clusters containing multiple novel enhancers driving *Chd7* expression in developing chick embryos. We validate enhancer activity *in vivo* and functionally determine key upstream transcription factors driving *Chd7* enhancers in the neural crest. Using chromatin accessibility analysis of donated human embryonic tissue, we find that neural crest-specific *CHD7* enhancers are highly conserved. Furthermore, we demonstrate that human enhancers are active in developing chick embryos, suggesting that the tissue-specific regulatory mechanisms enhancing *Chd7* in the neural crest are conserved between chicken and human. Our findings provide a potential mechanism to explain the aetiology of CHARGE syndrome where the *CHD7* gene is unperturbed. Finally, we demonstrate, for the first time, the upstream regulation of a ubiquitous chromatin-remodeller by tissue-specific transcription factors as an important mechanism to ensure enhanced chromatin remodelling activity in the developing neural crest.

## Results

*Chd7* was previously shown to be enriched in RNA-seq data from chick cranial neural crest cells (34) (Fig. S1A), where it was co-expressed with other neural crest genes (Fig. S1B) and clustered with pre-migratory neural crest markers (Fig. S1C). Here, we used fluorescent *in situ* hybridisation (Hybridisation Chain Reaction, HCR) (38) to resolve spatiotemporal *Chd7* expression in developing chick embryos. *Chd7* was detected at stage HH8 (39) within the cranial neural tube and pre-migratory neural crest cells as indicated by colocalisation with the neural crest marker *Sox10* and at low levels in the surrounding neuroectoderm (Fig. 1A). *Chd7* transcripts continued to overlap with *Sox10* in delaminating and migrating neural crest cells at HH9/10 and were also detected in the forebrain and neural tube at the vagal level from HH9 through HH15 (Fig. 1A). At later stages (HH13 - HH15) *Chd7* was more broadly expressed, with transcripts distributed across the head regions (midbrain and hindbrain) including the trigeminal ganglia (TG), developing face mesenchyme, eye and otic vesicle as well as a portion of vagal neural crest cells. In addition, at HH15, pharyngeal arches and dorsal root ganglion (DRG) cells were also *Chd7* positive (Fig. 1B).

**Fig. 1.**
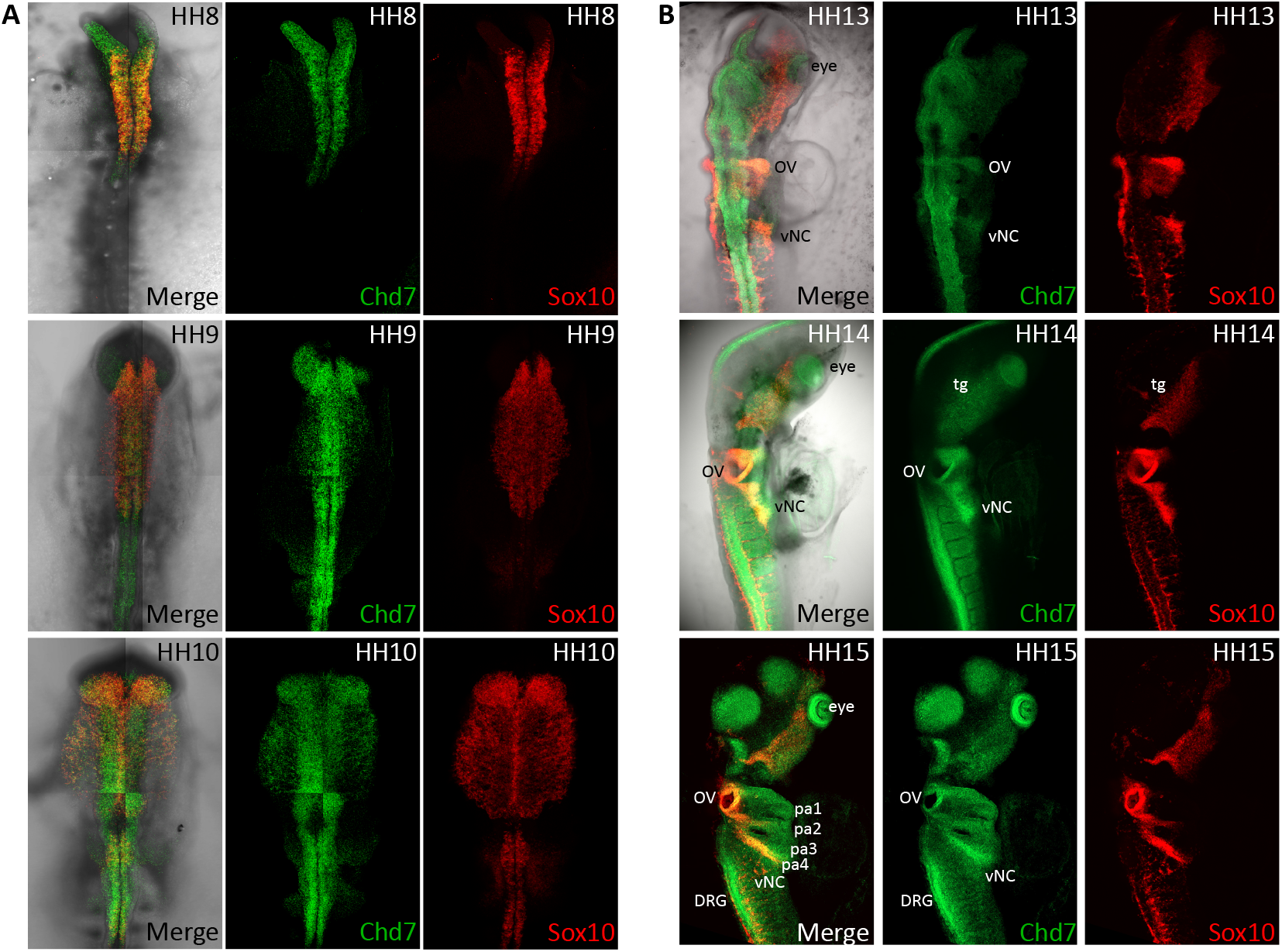
*Chd7* expression during early chick development. (A-B) *In vivo Chd7* expression (green) determined using HCR. *Sox10* (red) is used as a neural crest marker. (A) *Chd7* is expressed in pre-migratory and early migrating neural crest cells at HH8-10. (B) *Chd7* is more broadly expressed at HH13-HH15 including head and hindbrain structures. From HH13 through HH15 *Chd7* expression is detected in the otic vesicle (OV), trigeminal ganglia (tg) eye and vagal neural crest (vNC). At HH15 *Chd7* is also expressed in the pharyngeal arches 1-4 (pa1-4) and dorsal root ganglion (DRG).

### Epigenomic annotation of *Chd7* locus identifies numerous enhancer elements

*Chd7* was broadly expressed during early chick development, consistent with its omnipresent function as a chromatin remodeller, however, transcripts were notably enriched in the developing neural crest in-keeping with its known role in CHARGE syndrome. We next queried whether such a pervasive factor may be regulated in a tissue-specific fashion in the neural crest. To this end we explored neural crest-specific epigenomic features depicting putative enhancers interacting with the *Chd7* promoter. We first determined the *Chd7* topological associated domain (TAD) using Next-Generation Capture-C (40), a high-resolution targeted 3C approach adapted for low cell numbers. Using dissected dorsal neural tube tissue from HH8-10 chick embryos and differentiated red blood cells (RBC) from 10-day old chick embryos as a control, we resolved a broad neural crest specific TAD of approximately 1.1Mb, spanning ~85Kb upstream and ~1Mb downstream from the *Chd7* promoter, encompassing the entire gene body (Fig. 2A).

**Fig. 2.**
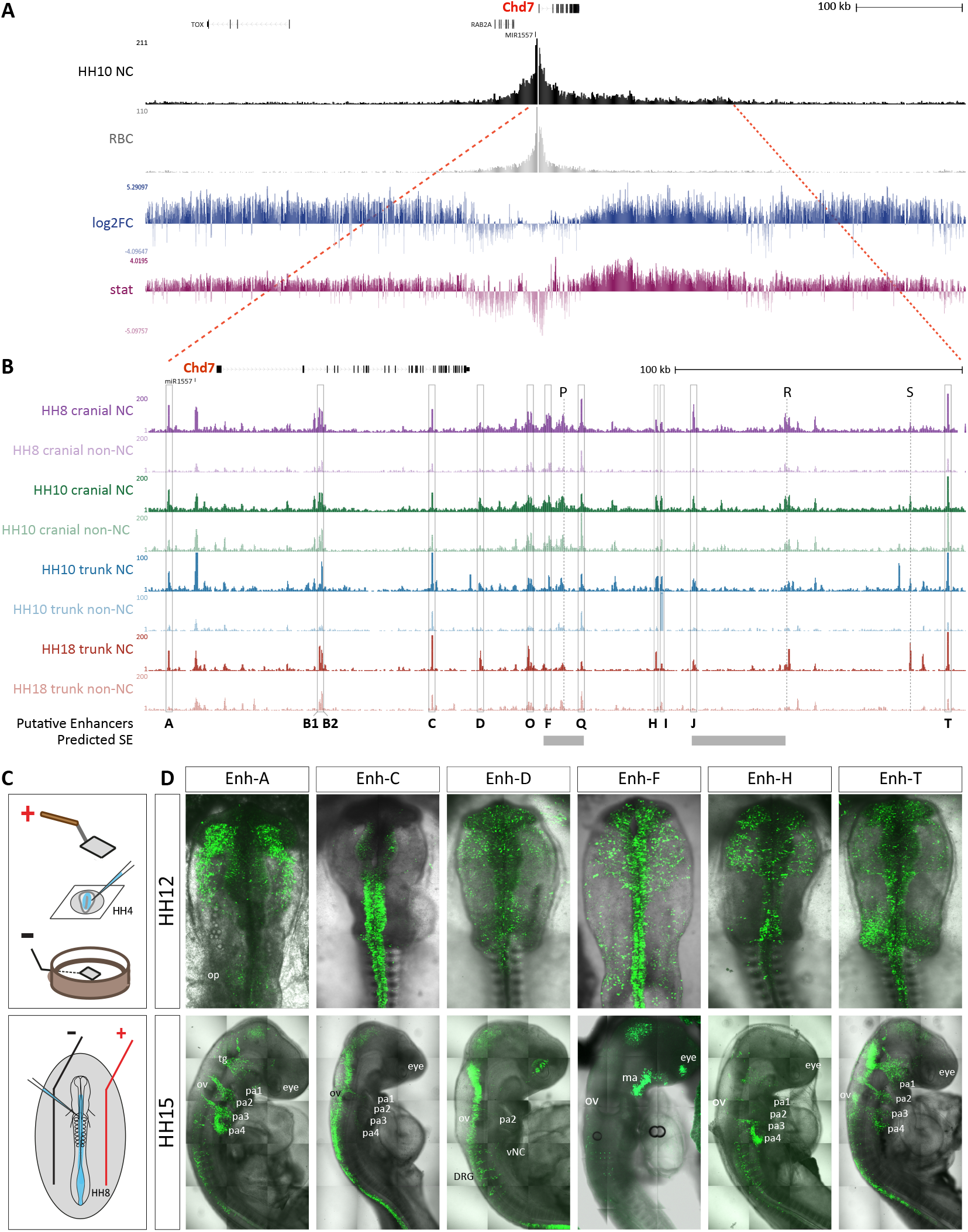
*Chd7* enhancer prediction from neural crest Capture-C and ATAC data. **(A/B)** UCSC genome browser view of the chick *Chd7* locus in galGal5. **(A)** Capture-C tracks from cranial neural crest at HH10 and control red blood cells (RBC), showing the *Chd7* TAD. Differential interactions were determined using DESeq2, hypothesis tested with Wald test and corrected for multiple testing using the Benjamin-Hochberg method. Wald statistics track (stat, in pink) represents ratio of LogFoldChange values and their standard errors. **(B)** ATAC-seq data from cranial (34) and vagal (41) neural crest cells and non-neural crest control cells collected by FACS using *FoxD3* enhancer NC1 (cranial) and *FoxD3/EdnrB* enhancers NC1/E1 respectively, at the time points indicated on figure. Putative enhancers within the *Chd7* TAD are indicated by grey boxes. Predicted super-enhancer (SE) regions are shown by grey bars. **(C)** Schematic representation of *ex ovo* and *in ovo* chick embryo electroporation techniques used to deliver enhancer reporter constructs. **(D)** Fluorescent reporter activity recorded from indicated enhancers at HH12 and HH15. pa; pharyngeal arch, tg; trigeminal ganglia, ov; otic vesicle, ma, mandibular arch.

We next surveyed the chromatin accessibility landscapes within the *Chd7* TAD using ATAC-seq data obtained from chick cranial (34) and vagal (41) neural crest and non-neural crest control cells at HH8-HH10 and HH10/HH18, respectively (Fig. 2B). We selected putative *Chd7* enhancers based on chromatin accessibility in neural crest cells. In total 14 putative enhancers were selected (A, B, C, D, F, H, I, J, O, P, Q, R, S, T) and screened for *in vivo* enhancer activity, using a fluorescent enhancer reporter assay (34, 42). In order to determine the full temporal range of enhancer activity we performed both *ex ovo* electroporation at HH4 and *in ovo* electroporation at HH8 (Fig. 2C) (43). These assays revealed 11 positive enhancers (Fig. 2D and Fig. S2) of which four were specifically active in neural crest cells (enh’s A, F, H, T) (Fig. 2D). Enh-A was the only upstream enhancer positioned approximately 18kb from the promoter. This element was accessible specifically in neural crest cells and not control environing tissues at both axial levels. At HH10-12, Enh-A enhancer activity was exclusively detected in delaminating and migrating cranial neural crest cells, (Fig. 2D and S2), while at HH15 enh-A activity was predominant in vagal neural crest cells migrating into the pharyngeal arches 2-4, as well as the head mesenchyme and trigeminal ganglia (Fig. 2D). Enh-B is located within the second intron of *Chd7*, ~37kb downstream from the promoter. Two large ATAC peaks (B1 and B2) were identified in the neural crest data, with smaller peaks resolved in the non-neural crest data, suggesting this region is accessible in a greater proportion of neural crest cells compared to non-NC cells. When tested *in vivo*, we found enh-B1 (+35.8kb from TSS) activity in a subset of pre-migratory neural crest cells at HH8, but present more broadly in neural tissue by HH12 (Fig. S2). Enh-B2 (+36.6kb from TSS) activity was restricted to the otic pla-code region at HH10-HH12 and later detected throughout the head and along the entire anteroposterior axis (HH18) (Fig. S2). Another intronic enhancer located between exons 22 and 23 (+75.3kb from promoter) (Fig. 2B), enh-C, was predominantly active in the hindbrain and trunk neural tube at HH12 and HH15, with some activity in the midbrain and migrating vagal neural crest cells (Fig. 2D), consistent with its accessibility profile (Fig. 2B). Downstream of the *Chd7* gene locus we identified 10 putative enhancers. Enh-D, +92.3kb from the TSS (~4kb) from the 3’UTR), opened progressively in the cranial neural crest data from HH8 to HH10, and in the vagal neural crest data from HH10 to HH18. Indicating an increase in the number of cells harbouring this element over developmental time (Fig. 2B). Enh-D presented broad activity in the forebrain and hindbrain regions including neural crest cells at HH12 (Fig. S2). Hindbrain activity continued at HH18 when reporter activity was also detected in the eyes, DRGs and vagal neural crest (Fig. S2). We previously identified a super-enhancer-like cluster of elements within the *Chd7* locus (*Chd7* SE) using H3K27ac and Brd4 ChIP data from migratory cranial neural crest cells (34) (Fig. S3). *Chd7* SE was located +114.5kb from the promoter, immediately downstream of enh-D and contained four distinct enhancers: enh-O, enh-F, enh-P and enh-Q (Fig. 2B). Enh-O (+109kb) activity was confined to the cranial neural tube at HH8-HH10 but was more broadly distributed across the cranial region including migrating neural crest cells at HH12. Later, (HH18) enh-O activity was also detected in the neural tube at the vagal level (Fig. S2), consistent with this enhancer’s accessibility in the vagal neural crest as well as non-neural crest cells at this stage (Fig. 2B). Enh-F (+115kb) activity was restricted to the cranial neural tube, including pre-migratory neural crest cells, at HH8. Enh-F activity persisted in delaminating and migrating neural crest cells at HH10 and HH12 (Fig. 2D and S2). At HH12, enh-F activity was also detected in the posterior cranial neural tube and some ectoderm cells particularly around the otic region (Fig. 2D). By HH15, enh-F activity was detected in head mesenchyme, eye and mandibular arch, but was absent from more posterior body structures consistent with the compaction of this element in vagal neural crest (Fig. 2B). At HH10-12 Enh-Q (+127kb) was active in a small number of cells in the otic placode region and later in the neural tube at vagal and cervical levels (Fig. S2). Genomic element termed Enh-P (+121kb) did not display any enhancer activity *in vivo*. We next tested two element lying within a 3kb region from each other, enh-H (+153kb) and enh-I (+155kb). Enh-H accessibility was consistently restricted to neural crest cells, while enh-H was active in the cranial neural crest at HH12 as well as in the neural tube at the level of rhombomeres 3-6 (Fig. 2D). At HH15, enh-H activity was recorded in the 3rd and 4th pharyngeal arch and the migrating trunk neural crest (Fig. 2D). Enh-I was accessible in neural crest cells and non-neural crest cells and was ubiquitously active at HH10-12 (Fig. S2). Enh-J (+166kb) was also ubiquitous in the head at HH12 but confined to face mesenchyme and trunk at HH15 (Fig. S2). Enh-T, the most distal downstream element tested (+255kb), was active in migrating neural crest cells as well as otic placode and vagal neural crest cells at HH12 (Fig. 2D). By HH15, enh-T activity was most prominent in pharyngeal arches, trigeminal ganglia and hindbrain, with lower activity levels in the neural tube and trunk neural crest. However, by this stage enh-T activity in the otic region was lost (Fig. 2D). Other putative elements tested (enh-R; +199kb and enh-S; +241kb (Fig. 2B) showed no enhancer activity.

By conducting a thorough survey of *Chd7* epigenomic landscape in the chick embryo, we revealed a large cohort of enhancers putatively controlling *Chd7* expression. Interestingly we found several enhancers specifically active in the neural crest and others displaying broader activity in other *Chd7*-positive tissues, thus demonstrating differential tissue-specific regulation.

### *Chd7* enhancers are conserved in human

While a large proportion of CHARGE syndrome cases are directly linked to mutations in *CHD7* coding region, there remains a significant number (>50%) of cases without causal annotation. We postulated that some instances of CHARGE syndrome may be indirectly linked to *CHD7*, via perturbation of upstream regulation of *CHD7* expression. To this end, we sought to identify regulatory elements controlling the expression of *CHD7* in human. We first examined epigenomic data obtained from *in vitro* derived human cranial neural crest cells (44). Specifically, we used ChIP-seq data for histone modifications associated with active enhancers (H3K4me1, H3K27ac), promoters (H3K4me3) and repressed chromatin (H3K27me3), as well as ATAC-seq data (Fig. 3A). We also generated ATAC-seq data from the dissected vagal region (otic placode to somite 7), cranial region and heart, as control, of a Carnegie Stage 11 (CS11) human embryo (approximately 28-30 days gestation). The sample was obtained from the Human Developmental Biology Resource, with consent from the donor for use in research of embryonic material arising from termination of the pregnancy. The homologues of chick *Chd7* enhancers were mapped in the human data using the LiftOver function from UCSC genome browser, with the exception of enh-A and enh-T, which were manually predicted based on synteny with other elements (Fig. 3A). We found that a number of elements identified in the human data as corresponding to chick *Chd7* enhancers, were marked by epigenomic features indicative of enhancer elements (Fig. 3A). We tested several of the human equivalent *CHD7* enhancers in the chick enhancer reporter assay. The chick neural crest specific enh-A was predicted to reside within the first intron of human *CHD7* gene. While the putative human enh-A element did not appear accessible in the *in* vitro-derived neural crest cells, this region was marked by H3K4me1, suggesting this was a poised element in human neural crest. This observation was corroborated by a significant accessibility (large peak) in the human embryonic ATAC-seq dataset. Human Enh-B was marked by H3K4me1 and H3K27ac, and the element was accessible in *in vitro*-derived neural crest cells and human embryonic vagal neural crest (Fig. 3A). *In vivo*, human enh-B was predominantly active in the developing forebrain region at HH12 and was also seen in the midbrain and neural tube extending into the trunk region (Fig. 3B, panel i). Compared to chick enh-B which was more predominant in the hindbrain and vagal neural tube but also active, albeit slightly weaker, in the forebrain region (Fig. 3B, panel ii). At HH15 human enh-B was detected along the neural tube and across the developing face (Fig. 3B, panel iii), similar to the chick element (Fig. S2). At HH18 we resolved somite specific activity from both chick and human enh-B (Fig. 3B, panels iv and v). Human enh-C had no enhancer associated features in the *in vitro*-derived neural crest data. However, since this data was generated from cranial neural crest cells (44) and enh-C was primarily active in the neural tube at the trunk level in chick this is unsurprising and indeed reiterates the tissue specificity of *Chd7* regulation. Human enh-C was active in the trunk neural tube at HH12 which was maintained at HH15 where we also detected enhancer activity in the pharyngeal arches (Fig. 3C, panel ii inset). Co-electroporation of chick and human enh-C showed the human element very precisely recapitulated the pattern of chick enh-C in the trunk neural tube at HH12 (Fig. 3C panels iii, iv, v). Enh-D and enh-Q were marked by H3K4me1, but showed no accessibility in the *in vitro*-derived neural crest cells, whereas enh-Q was accessible as per the human embryo data (Fig. 3A). Enh-T was also characterised by H3K4me1 modification and found accessible in both data sets; this region was also marked by H3K27me3 possibly indicative of a repressive state. Enh-O showed some accessibility in the human data sets and was marked by H3K4me1 and H3K27ac with some indication of H3K4me3 suggesting this is indeed an active enhancer in human neural crest. Enh-F was accessible in the human embryonic neural crest and whilst the region was decorated by H3K4me1, significant H3K27me3 signal was also present, again indicating repressor activity. Enh-H and enh-I could not be distinguished from each other, but this region was accessible and marked by H3K4me1.

**Fig. 3.**
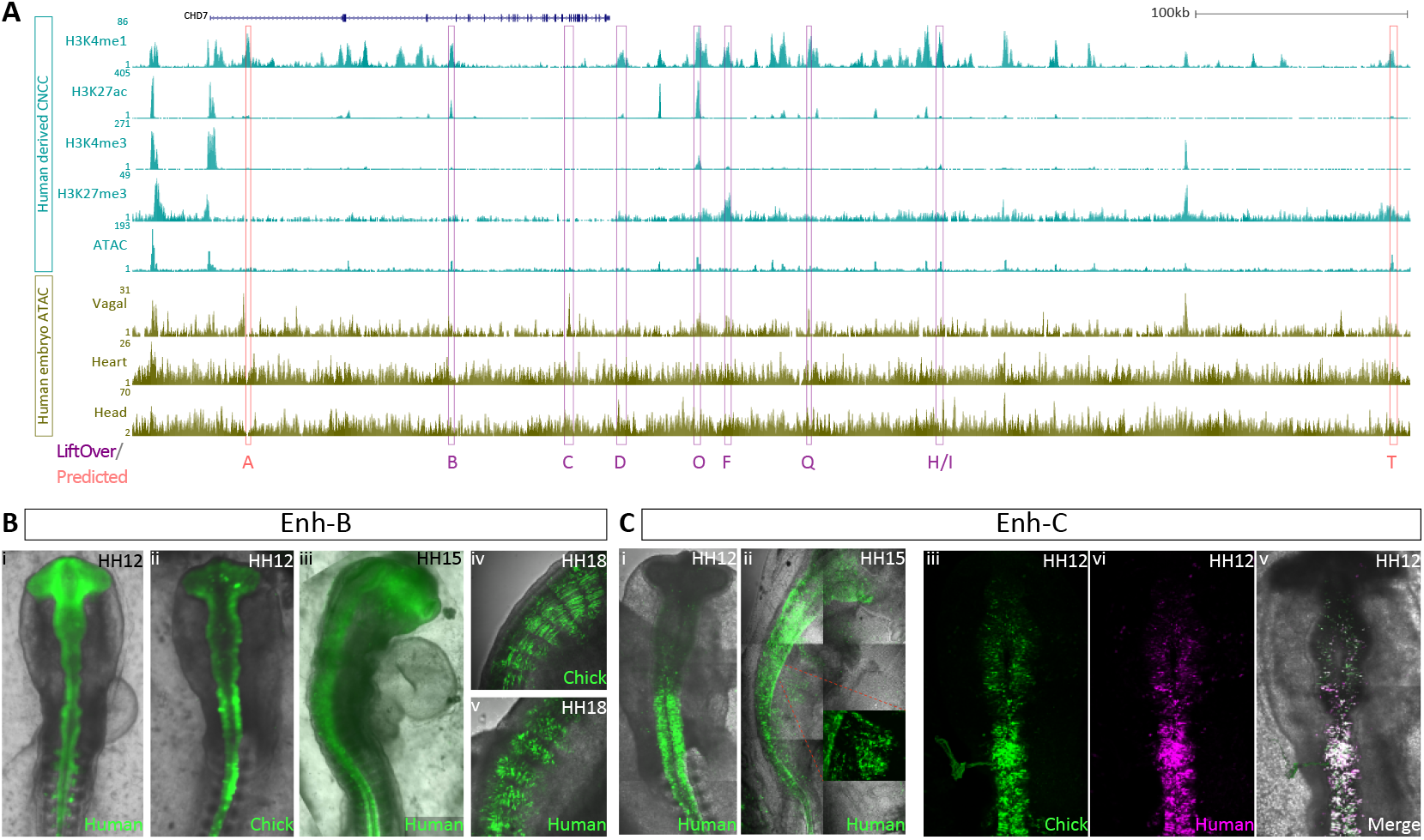
Epigenomic landscape of human *CHD7* locus. **(A)** *CHD7* locus in human genome (hg38), showing ATAC-seq and histone ChIP-seq data from human *in vitro* derived cranial neural crest cells (44) and ATAC-seq data from vagal, cranial and heart tissue of a human embryo (CS11). Relative positions of chick enhancers were detected using the UCSC LiftOver function or predicted by synteny. (B) Human enh-B activity *in vivo* at HH12 (i), HH15 (iii) and HH18 (v) compared with chick Enh-B activity (ii and iv). (C) Human enh-C activity atHH12(i) and HH15 (ii). Co-electroporation of Human Enh-C (magenta) and chick enh-C (citrine) at HH12 shows co-localisation of enhancer activity (iii – v).

By comparing human and chick loci, we demonstrate conserved cis-regulation of *CHD7* at least across amniotes. Our findings illustrated the utility of *in vitro* derived data, whilst highlighting important differences observed in the *in vivo* embryonic context. Furthermore, this provides an alternative mechanism by which *CHD7* expression can be affected in CHARGE patients, where mutations are not detected in the gene itself.

### *Chd7* enhancers are differentially regulated by neural crest transcription factors

Enhancer activity is determined by the combinatorial binding of transcription factors (TF) to their cognate motifs within the enhancer sequence. Indeed, such interactions are the foundation for cell-type specific regulation of gene expression. In order to explore putative TF binding within the *Chd7* enhancers, we first scanned human TF binding motifs, allowing us to identify motif instances within each enhancer sequence (Fig. S4). Next, we calculated the enrichment of binding sites within each enhancer using the log odds ratio score (Fig. 4A). The two approaches yielded broadly similar results. Across all *Chd7* enhancers we found an enrichment of neural crest associated TF’s, such as Tfap, SoxE (Sox8/9/10) and SoxB1 (Sox2/3) factors as well as a significant input from the Zic family members (Fig. 4A). Many enhancers received input from Arnt/Arnt2 and Atf2 factors, which we previously described in our survey of core upstream TFs driving neural crest lineages (34). Given the broad spatial and temporal range of *Chd7* enhancers, it is likely that different TF combinations drive enhancer activity at different time points and in different locations. In particular, we noted that the neural crest specific enhancers enh-A and enh-T were differentially enriched for Tfap2 and SoxE factors respectively (Fig. 4A, S4). Suggesting differential mechanisms for *Chd7* regulation within the same cell population. Enh-T was driven by the Tfap2 family members which were not predicted to bind and drive enh-A. Enh-T was the only enhancer enriched for Sox8 binding, in addition to Sox10 and Sox2/3 factors (Fig. 4A). Given the activity of enh-T extends to the otic placode region (Fig. 2D) and Sox10, Sox8 and Sox3 are all expressed here (45,46) it is likely these transcriptional activators and repressors work to balance enh-T activity in the otic placode region. Conversely, Enh-A was enriched for Sox9 and Sox10, but not Sox8 binding with presence of repressive Sox2/3 sites, (Fig. 4A), likely repressing enh-A activity in the otic region (Fig. 2D). Enh-H shared a similar signature with enh-A, whereby both enhancers lack Tfap2 sites but are driven by Sox9/10 and Sox2/3. Enh-A is also repressed by Snai2 in the hindbrain neural tube region. Enh-F and enh-C were also enriched for Snai2 binding (Fig. 4A) but were active in the hindbrain neural tube (Fig. 2D), likely driven by Pax6/7 which is notably absent in enh-A (Fig. 4A). Enh-T and enh-H did not contain Snai2 repressor sites (Fig. 4A), allowing enhancer activity in neural crest cells and the otic placode region (Fig. 2D). Enh-F, which is also active in the cranial neural crest was enriched for both SoxE/B and Tfap2 factors. Furthermore, both enh-F and enh-C contained a Pax6/7 motifs (Fig. 4A), and both enhancers were active in the neural tube from the hindbrain region and more posteriorly at HH12 (Fig. 2D). At HH15 enh-F is active in the developing eye (Fig. 2D) consistent with Pax6 expression whereas enh-C activity continues in the trunk region (Fig. 2D) consistent with Pax7 expression. Plus shared Pax6/7 motif likely represents differential regulation of these enhancers by related factors over different spatio-temporal contexts.

**Fig. 4.**
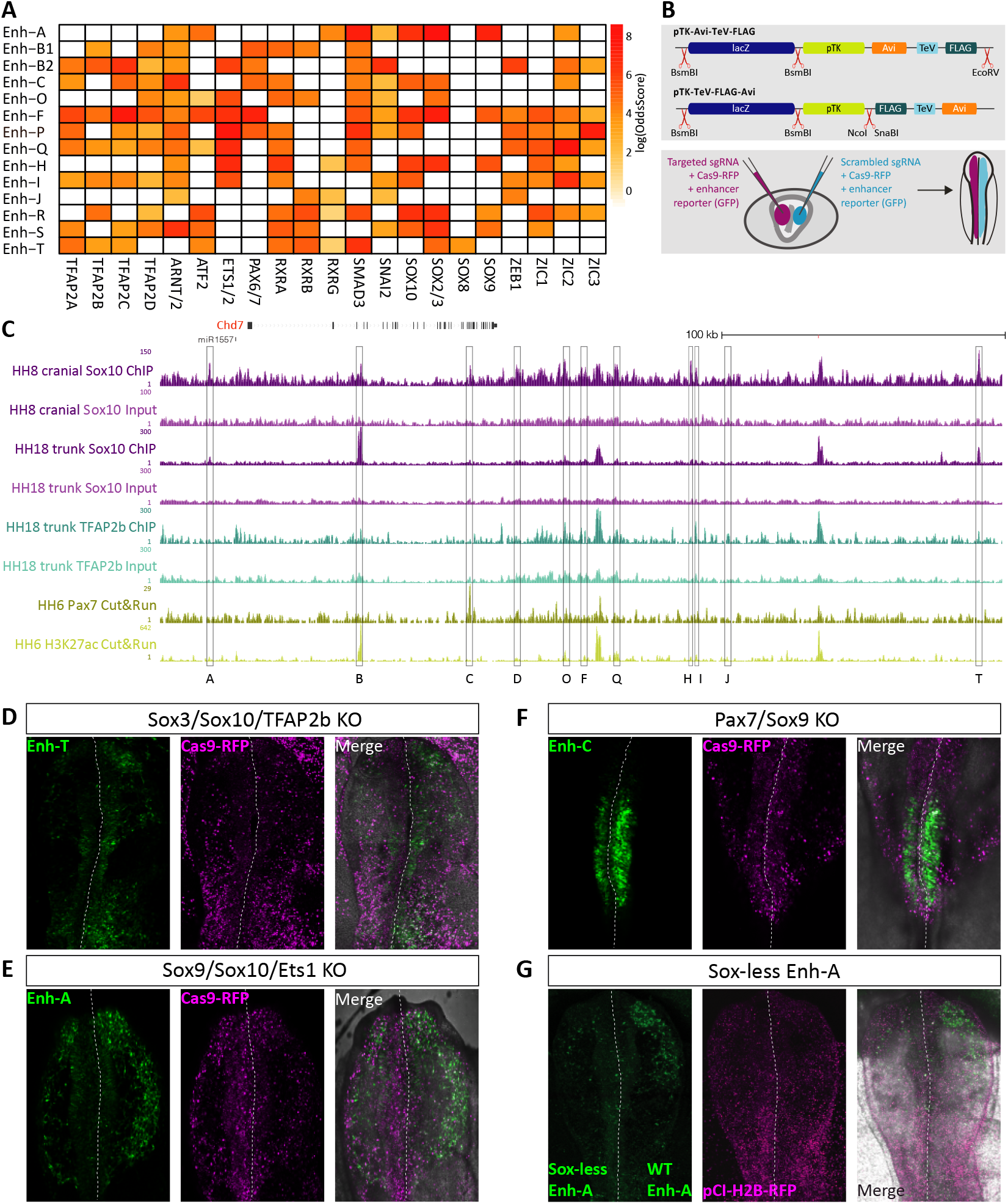
Upstream transcription factor signatures driving *Chd7* enhancer activity. (A) Heatmap showing predicted TF binding motifs within chick *Chd7* enhancers. (B) Schematics depicting biotin-ChIP assay (top panel) and bilateral electroporation (bottom panel). (C) Tracks from UCSC genome browser showing biotin-ChIP data for Sox10 (purple), Tfap2B (teal) and Pax7 (yellow). (D-F) Chick embryos showing indicated enhancer activity (GFP) and Cas9 (RFP) following bilateral electroporation of Cas9 and guide RNAs targeting selected TFs on the left (experimental) and scrambled guide RNA with Cas9 on the right (control). (G) Chick embryo bilaterally electroporated with a construct containing enh-A where all the SoxE sites have been mutated on the left, and wild type enh-A construct on the right, both driving GFP expression. pCI-H2B-RFP was co-electroporated on both sides as a control.

Whilst broadly present, Zic factors were largely absent from neural crest enhancers (enh-A, T, C). Only enh-F, active in early cranial neural crest cells as well as some ectoderm cells in the hindbrain region, contained binding sites for all Zic factors (Zic1, 2, 3), suggesting this family regulates non-neural crest enhancer activity, consistent with the broad expression patterns of these factors in the neural tube (47). Motif analysis suggests another canonical neural crest regulator, Ets1, also plays a role in *Chd7* regulation. We detected an enrichment of Ets1 binding sites in enh-H, enh-F (Fig. 4A) and also identified some sites in enh-A (Fig. S4).

Given previously established relationships between Chd7 and Sox10 (48, 49) we set out to explore the possibility that Sox10 could also be acting upstream of *Chd7*. We profiled Sox10 binding genome-wide using Biotin-Streptavidin based Chromatin Immuno-Precipitation (biotinChIP). For this purpose, we generated constructs to facilitate Avi-tagging of transcription factors at the N- or C-terminal. Furthermore, we incorporated an enhancer cloning site upstream of the expression cassette, such that the Avi-tagged-TF would be tissue specifically expressed under the control of its cognate enhancer (Fig. 4B) (Addgene #110204, #110205, see methods). We used the *Sox10* enh-99 (34) to drive C-terminally tagged Sox10. Following co-electroporation with ubiquitously expressed biotin ligase, BirA, (Addgene #127781) (41) into HH4 embryos the Avi-tagged-TF is biotinylated *in vivo*, facilitating highly stringent isolation using streptavidin beads (Fig. 4B) (50). This approach allowed us to isolate specific TFs of interest at different stages and examine bound regions genome-wide. All active *Chd7* enhancers showed some binding by Sox10, compared to the ChIP input sample (Fig. 4C), indicating Sox10 acted as a key player in *Chd7* regulation. Neural crest specific enhancers, enh-A and enh-T showed particularly high signal for Sox10 binding, consistent with our motif analysis (Fig. 4A, S4). Enh-H and enh-F were also significantly bound by Sox10, driving their neural crest-specific activity. Other enhancers whose activity was not restricted to the neural crest including enh-B, enh-C, enh-D and enh-O were also bound by Sox10 suggesting Sox10 drives their activity in neural crest cells with other factors driving their activity in non-neural crest tissues.

Given the observed enrichment of Tfap2B motifs (Fig. 4A) we next examined Tfap2B binding at *Chd7* enhancers, Tfap2B is a known neural crest specification factor. Using Tfap2B biotin-ChIP data generated from cranial and vagal neural crest of HH18 embryos, where Tfap2B-Avi was driven under the control of the Ednrb enhancer, E2, (Addgene #127776) (41). This data confirmed Tfap2B indeed binds to enh-T, enh-F, enh-B as well as enh-Q and enh-I as predicted by our motif analysis (Fig. 4A, S4). Furthermore we did not detect Tfap2B binding at enh-A, consistent with the absence of enrichment in Tfap2 motifs at this enhancer (Fig. 4A). Using the Cut&Run (Cleavage Under Targets and Release Using Nuclease) assay (51, 52), we next profiled Pax7 binding in dorsal neural tube tissue extracted at HH8-9. Here we confirmed binding of Pax7 at enh-C, but did not detect Pax7 binding at enh-F (Fig. 4A), supporting the hypothesis that Pax6 occupies this motif in enh-F.

### Functional perturbation of upstream transcription factors disrupts enhancer activity

To further resolve key TF combinations required for *Chd7* enhancer activity we used the chick CRISPR/Cas9 system (53) to disrupt multiple TFs simultaneously. By co-electroporating enhancer reporter constructs with target guide RNAs (gRNAs) and Cas9 plasmids, in a bilateral fashion (Fig. 4B), we were able to directly assess the effects of TF loss on enhancer activity (Fig. 4D-G).

As the predominant TF bound by all *Chd7* enhancers and a neural crest master regulator, we primarily considered knock-out combinations including *Sox10*. We also focused our functional assessments on the neural crest specific enhancers, enh-A, enh-T and enh-C. As described above, enh-T was bound by Sox10 (Fig. 4C) in addition this enhancer was enriched for the Tfap2B and Sox2/3 motifs (Fig. 4A). Similarity of familial motifs can make it difficult to discern the specific factor occupying a particular site, however given enh-T is also active in the otic placode region, where *Sox3* but not *Sox2* is expressed, we hypothesised that this motif represented Sox3 binding. Using gRNAs previously described (34, 41) we targeted *Sox10* in combination with *Sox3* and *Tfap2B* on the left side of the embryo and applied a control, scrambled gRNA to the right side. Both sides received Cas9-RFP and enh-T driving Citrine (Fig. 4D). Concomitant *Sox10/Sox3/Tfap2* knockout resulted in a reduction of enh-T-driven Citrine expression on the left side of the embryo, compared to the control right side, in the migrating neural crest cells, neural tube and the otic region (100% embryos, n=10) (Fig. 4D).

Enh-C was the only *Chd7* enhancer with predominant activity in the trunk neural tube (Fig. 2D). TF motif analysis revealed enh-C was enriched for Pax7 motifs (Fig. 4A) and also contained motifs for Sox9 binding (Fig. S4), while both *Pax7* and *Sox9* are expressed in the trunk neural tube. Combined knock-out of *Pax7* and *Sox9* caused a reduction in enh-C activity specifically in the trunk neural tube at HH10 (Fig. 4E). We next sought to determine crucial TF combinations driving enh-A activity. Sox9 and Sox10 sites were both enriched in this enhancer (Fig. 4A). Considering SoxE factors are highly expressed in the neural crest, share similar motifs and frequently collaborate we reasoned these could be core upstream factors for enh-A activity. We also identified Ets1 as a putative upstream factor (Fig. S4). Simultaneous knock-out of *Sox9, Sox10* and *Ets1* caused a profound reduction in enh-A activity in the migrating neural crest at 10ss (Fig. 4F). Sox2/3 sites were also enriched in enh-A however, combined knock-out of these factors had no effect on enh-A driven Citrine expression, suggesting these factors act as canonical transcriptional repressors to inhibit enhancer activity in non-neural crest cells, consistent with their expression in neural tissue and otic placode. In parallel, we generated a mutant construct of enh-A whereby all the SoxE sites were mutated this resulted in reduced enhancer function (Fig. 4G) reinforcing the notion that enh-A is primarily driven by SoxE factors.

Using functional perturbation experiments we have corroborated our motif analysis and ChIP data, demonstrating that *Chd7* neural crest enhancers are indeed functionally dependent on neural crest transcription factors.

## Discussion

Transcriptional regulation of gene expression is mediated by cis-regulatory elements (enhancer and promoters) that integrate the action of upstream transcription factors upon the downstream target gene(s). Developmental and housekeeping enhancers differ in their genomic location and motif complement, which in turn dictates their interaction dynamics with core promoter sequences (54). Developmental genes are regulated by distally located enhancers containing celltype specific transcription factor motifs facilitating spatial and temporal control of gene expression. Whereas expression of housekeeping genes is mediated by TSS proximal enhancers containing a different profile of ubiquitous motifs enabling stable core promoter interactions. Reports of tissue specific regulation of housekeeping genes by distal acting enhancers are rare.

Despite its broad expression and housekeeping function, heterozygous loss of *CHD7* leads to CHARGE syndrome in humans; a tissue-specific neurocristopathy affecting structures patterned by, or derived from, neural crest cells. Tissue specific function of housekeeping genes has previously been attributed to downstream interactions with tissue specific transcription factors. For example, in Schwann cells, the transcription factor Sox10 recruits chromatin-remodelling complexes to cis-regulatory regions of *Oct6* and *Krox20* genes (55), whereas in Oligodendrocytes, Olig2 is reported to direct Smarca4/Brg1 to *cis*-regulatory regions of genes which control their differentiation (56). Similarly, tissue-specific transcription factors were reported to directly recruit and target PRC1 to chromatin in megakaryocytic cells (57). Furthermore, previous CHD7 studies resolved downstream targets and partners for CHD7 activity, for example, Chd7 is reported to bind to enhancers of the neural crest transcription factor *FoxD3* in mouse neural crest stem cells along with Oct3/4, Sox2 and Nanog (32). And in human neural crest cells CHD7 binds with PBAF at a distal regulatory element upstream of *SOX9* (29).

Here, we probed the intriguing possibility that neural crest enrichment of a housekeeping factor such as *Chd7*, may be regulated by tissue-specific *cis*-regulatory elements and their upstream transcription factors. In doing so, we also established the position of *Chd7* in the neural crest gene regulatory network.

Consistent with previous work in the chick (58), we saw *Chd7* expression initiated at HH8 along the cranial neural tube including pre-migratory neural crest cells. *Chd7* was upregulated in neural crest cells throughout their emergence from the dorsal neural tube and during their subsequent migration. *Chd7* transcripts were also detected in placode derivatives including the eye and otic vesicles, in addition to the pharyngeal arches, dorsal root ganglia and trigeminal ganglia. This pattern of *Chd7* expression is conserved in mice (28), Zebrafish (59) and Xenopus (29).

Using regulatory data from isolated cranial pre-migratory and early migrating chick neural crest cells, we identified a group of eleven novel *Chd7* enhancers. *In vivo* activity of which collectively recapitulated *Chd7* expression. While some enhancers display specific activity within the cranial neural crest, others had broader activity in the head and/or trunk and vagal neural crest. We found enhancers enh-A, enh-H and enh-T were also active in the migrating cardiac neural crest, however we did not observe any enhancer activity in the developing heart itself, consistent with the lack of *Chd7* expression here at the stages studied. It is likely that heart specific cis-regulation initiates at later stages when *Chd7* is turned on in this tissue (60). Indeed, despite the high background, we could depict some heart specific chromatin accessibility in our human embryo ATAC-seq data. This would suggest *CHD7* expression is regulated by different factors within sub-populations of neural crest cells. Importantly, we found conservation of a number of the *Chd7* chicken enhancers in epigenomic data obtained from human *in vitro* derived cranial neural crest cells and ATAC-seq data from E9.5 human vagal tissue. Human enh-B and enh-C were both active in the chick embryo where reporter activity overlapped with that of their chicken counterparts. Iterating the conservation of *Chd7* regulation during embryo development, and indeed supporting the utility of the chick model for exploring human conditions. While many clinical CHARGE syndrome cases contain mutations in the *CHD7* Open Reading Frame (ORF), it is possible that the remaining unattributed cases are caused by disruptions in cis-regulatory elements controlling *CHD7*. Thus, the enhancers identified here represent potential alternative screening sites for CHARGE syndrome patients without mutations within the *CHD7* gene body.

Enhanceropathies are a group of clinical conditions caused by mutation or misregulation of enhancers. This can occur via direct mutation of the enhancer sequences, e.g. the point mutation in a sonic hedgehog enhancer that causes polydactyly (61), or mutation of upstream genes that produce proteins which regulate enhancers e.g. Kabuki syndrome (62). Interestingly, Kabuki syndrome is attributed to mutations in members of the polycomb repressive complex-2 (PRC2) (63) and has overlapping features with CHARGE syndrome (64). It is therefore important to understand which motifs within an enhancer are required for transcription factor binding to induce enhancer activity. As such, we used transcription factor motif predictions to identify the core factors responsible for *Chd7* enhancer activity. In doing so we reveal potentially sensitive sites for disease causing mutations. We validated motif predictions using ChIP assays and found the neural crest master regulator Sox10 binding at neural crest specific enhancers enh-A and enh-T. Functional relationships between Sox10 and Chd7 have been previously described. *Sox10* is dysregulated in *Chd7* morphant zebrafish (48) and co-IP experiments have shown Sox10 and Chd7 physically interact to regulate downstream targets (49). Here, we extend this relationship by demonstrating that Sox10 also acts as an upstream driver for *Chd7* expression within the neural crest by regulating enh-A and enh-T. We also identified Tfap2B binding at enh-T. TFAP2B and CHD7 are known to interact in neural progenitor cells (65), but similarly to *SOX10, TFAP2B* is reported to be reduced upon loss of *CHD7* in that context. Our work demonstrates that Tfap2B is also driving *Chd7* enhancer activity in neural crest cells, potentially indicating the existence of positive feedback loops between Sox10/Tfap2B and Chd7. We also validated Pax7 binding enh-C, which is active throughout the trunk. This is the first report of Pax7 regulating *Chd7*, and provides evidence of axial specific regulation of *Chd7*. Furthermore enh-C has a conserved and active counterpart in human, moreover, mutations in *PAX7* have recently been shown to affect craniofacial development (66), a phenotypic feature of CHARGE syndrome. Our data therefore offers an additional mechanism by which PAX7 functions to regulate normal neural crest ontology. Curiously, a significant proportion of *Chd7* enhancers contained retinoic acid receptor sites, RXRA has previously been described as a Chd7 interacting protein that is mutated in congenital heart disorders (60). We also found a subset of *Chd7* enhancers were enriched for Zic binding sites. ZIC1/4 are responsible for cerebellar vermis anomalies in Dandy-Walker syndrome (67), which is also a phenotype of CHARGE syndrome. *Zic1/2* gene-deficient mice share anterior hemisphere foliation defects with *Chd7* deficient mice (68). This demonstrates the overlap of phenotypes shared between neurocristopathies and other developmental defects and illustrates the need for more mechanistic understanding of the complex genetic relationships underlying such conditions, to improve diagnosis and clinical management.

In summary, here we provide the first report of upstream tissue specific regulation of the chromatin remodeller, *Chd7*, during early chick neural crest development. We detail enhancer activity and cognate transcription factor binding, embedding *Chd7* within the neural crest gene regulatory network, downstream of neural crest master regulators including Sox10, Tfap2B and Pax7. Thus, providing a critical link between neural crest transcription factors and *Chd7* activity, and ultimately providing a putative mechanism as to how they effectuate normal development of neural crest derived tissues. Furthermore, we demonstrate features of *Chd7* regulation are conserved in human, offering a potential resource for identifying causative mutations driving CHARGE syndrome as well as a novel mechanism for tissue specific activity of omnipresent factors.

## ACKNOWLEDGEMENTS

We thank all members of the Sauka-Spengler lab for helpful discussions and feedback. High-resolution imaging was conducted within the Wolfson Imaging Centre at the MRC WIMM.

## Materials and Methods

### Chicken embryo collection

Chick embryos were harvested from fertilised Bovans Brown chicken eggs (Henry Stewart & Co) were incubated at 37°C with approximately 40% humidity. Embryos were staged according to the Hamburger and Hamilton table of normal chick development (39). All experiments were performed on chicken embryos younger than 12 days of development, and as such were not regulated by the Animals (Scientific Procedures) Act 1986.

### Embryo preparation and electroporation

For *ex ovo* electroporation, gastrula stage (HH4) embryos were captured using the filter paper based ‘easy-culture’. Plasmids were injected across the whole the epiblast and electroporated with 5 pulses of 5V, 50ms on, 100ms off. Bilateral electroporation was used for perturbation experiments, whereby control and experimental regents are delivered to opposite sides of the primitive streak, providing ideal internal, stage matched controls for each experiment. Embryos were cultured on albumin at 37°C 5% CO_2_ overnight to the desired stage. For full procedure see (43). For *in ovo* electroporation, eggs were incubated on their sides until HH8/9. A window was made in the egg and plasmids were injected into the lumen of the neural tube along the whole anterior posterior axis. Electroporation was at conducted at 3 pulses of 12.5V, 50ms on, 100ms off. To ensure electroporation of plasmids across both sides of the neural tube the polarity was inverted and a second set of pulses was applied with the same settings.

### Enhancer cloning and testing

Putative enhancer elements were cloned and tested individually as described in detail here (42).

### Fluorescent *in situ* hybridisation chain reaction (HCR)

Fluorescent *in situ* hybridsation chain reaction was performed using the v3 protocol (38). Briefly, embryos were fixed in 4% paraformaldehyde (PFA) for 1 hour at room temperature, dehydrated in a methanol series and stored at −20°C overnight. Following rehydration embryos were treated with Proteinase-K (20mg/mL) for 1-2.5 min depending on stage (1 min HH4-6, 2.5 mins for older embryos) at room temperature and post-fixed with 4% PFA for 20 min at room temperature. Embryos were washed in PBST for 2x 5 min on ice, then 50% PBST / 50% 5X SSCT (5X sodium chloride sodium citrate, 0.1% Tween-20) for 5 min on ice and 5X SSCT alone on ice for 5 min. Embryos were then pre-hybridised in hybridisation buffer for 5 min on ice, then for 30 min at 37°C in fresh hybridisation buffer. Probes were prepared at 4pmol/mL (in hybridisation buffer), pre-hybridisation buffer was replaced with probe mixture and embryos were incubated overnight at 37°C with gentle nutation. Excess probes were removed and embryos washed with probe wash buffer for 4x 15 min at 37°C. Embryos were pre-amplified in amplification buffer for 5 min at room temperature. Hairpins were prepared by snap-cooling 30pmol (10ml of 3mM stock hairpin) individually at 95°C for 90 sec and cooled to room temperature for minimum 30 min, protected from light. Cooled hairpins were added to 500μl room temperature amplification buffer. Preamplification buffer was removed from embryos and hairpin solution was added overnight at room temperature, protected from light. Excess hairpins were removed by washing in 5X SSCT 2x 5 min, 2x 30 min and 1x 5 min at room temperature.

### Imaging analysis

Enhancer electroporated or HCR processed embryos were were mounted on slides and imaged using Zeiss LSM 780 Upright confocal microscope. Images were processed using Zeiss Zen software, Z-stacks scans were collected at 6μm intervals across approximately 70-100μm, maximum intensity projections of embryo z-stacks are presented. Tile scanning was used and stitched using bidirectional stitching mode, with overlap of 10%.

### Human embryo ATAC-seq

The CS11 embryo was provided by the Human Developmental Biology Resource (HDBR - https://www.hdbr.org/general-information). HDBR has approval from the UK National Research Ethics Service (London Fulham Research Ethics Committee (18/LO/0822) and the Newcastle and North Tyneside NHS Health Authority Joint Ethics Committee (08/H0906/21+5)) to function as a Research Tissue Bank for registered projects. The HDBR is monitored by The Human Tissue Authority (HTA) for compliance with the Human Tissue Act (HTA; 2004). This work was done as part of project #200295 registered with the HDBR. The material was collected after appropriate informed written consent from the donor by medical termination. The sample was collected and transported in cold L15 media. It was then transferred to M2 media and imaged on a Leica Stereo microscope prior to processing. It was karyotypically normal and female ((rsa(13,15,16,18,21,22, X)x2). The sample was dissociated to a single-cell solution and ATAC-seq was conducted as previously published (69).

### Biotin ChIP-seq

Avi-tagged transcription factor constructs (1.0μg/μl) were co-electroporated with a pCI NLS-BirA-2A-mCherry plasmid (0.5μg/μl) into HH4 embryos (ex ovo) for collection at HH8 (Sox10) or HH8/9 (in ovo) for collection at HH18 (Tfap2B). The Biotin ChIP-seq procedure is described in detail here (50).

### Genome editing

Guide RNAs were designed to target genes and co-electroporated with pCI-Cas9-Citrine (Addgene #92358. For more detail see (53).

### Cut and Run

Cut and Run was performed according to published protocols (51, 52). **Cell preparation:** Embryos (x10 per stage) were collected at the desired stage and placed in 1ml nuclei extraction buffer (0.5% NP40, 0.25% Triton-X, 10mM Tris-HCl-pH 7.5, 3mM CaCl_2_, 0.25M sucrose, 1mM DTT, 0.2mM PMSF, 1X protease inhibitor tablet), following 10 strokes in a glass dounce homogeniser with pestle A on ice, the solution was centrifuged 600g for 3 min at room temperature in a low-binding 1.7ml Eppendorf tube. Nuclei extraction buffer was removed and cells were washed twice with DIG-wash buffer (20mM HEPES pH 7.5, 150mM NaCl, 0.5mM Spermidine, 0.05% digitonin, 1X protease inhibitor tablet). **Binding cells to beads and antibody incubation:** Pelleted cells were then resuspended in 1ml DIG-wash buffer and Concanavalin-A beads were added while shaking in an Eppendorf ThermoMixer at moderate speed. After 10 min rotating, the bead-cell solution was divided into aliquots, one for each antibody to be used. The beads were collected on a magnetic stand, DIG-wash buffer was removed and 150μl of antibody buffer (2mM EDTA in DIG-wash buffer) was added to each aliquot while shaking. Primary antibody was added at 1:100 dilution (αChicken Pax7 (DSHB), H3K27ac Abcam Ab4729) and samples were incubated overnight at 4°C with end-over-end rotation. **Binding Protein-A/G-MNase fusion protein:** Following a brief spin, beads were collected on a magnetic stand and washed twice with 1ml DIG-wash buffer. While shaking, 150μl of the Protein-A/G-MNase solution was added to the beads and incubated for 1 hour at 4°C with rotating. Beads were collected on the magnetic stand and washed twice with DIG-wash buffer and resuspended in 100μl DIG-wash buffer while shaking. Tubes were transferred to a cold block on ice, chilled to 4°C, 2μl of 200mM CaCl_2_ was added with shaking and promptly placed back in the cold block on ice for 30 min. While shaking 100μl of STOP-buffer (340mM NaCl, 20mM EDTA, 4mM EGTA, 0.05% Digitonin, 100μg/ml RNase A, 50μg/ml Glycogen) was added and incubated at 37°C for 30 min. **Collection of fragmented antibody bound DNA:** Beads were collected on a magnetic stand and the supernatant, now containing digested chromatin, was transferred to a new 1.7ml low-binding Eppendorf tube. Chromatin proteins were degraded by adding 2μl of 10% SDS, 2.5μl of Proteinase-K (20mg/ml) and incubated at 50°C for 1 hour. DNA fragments were cleaned up using standard phenol:chloroform extraction and ethanol precipitation and finally resuspended in 20μl water. Following quality control measures on Agilent Tapestation (high sensitivity) and Qubit, sequencing ready libraries were made using the NEBNext^®^ Ultra™ DNA Library Prep kit and sequenced on the Illumina Next-seq 500/550 platform using High Output Kit v2 (75 cycles).

## Data analysis

### ATAC-seq

Sequencing files were demultiplexed and the resulting files were merged. Read quality evaluation was performed using FastQC v0.11.4 (70). Nextera adaptor sequences were trimmed using TrimGalore (v0.4.1 settings: -nex-tera-paired-three_prime_clip_R1 1-three_prime_clip_R2) and then mapped to the human genome Hg38 assembly using bowtie (v1.0.0 settings: -S -X 2000) (71). Only aligned pairs were retained based on binary alignment map (BAM) flags, and mitochondrial reads were removed from the BAM file using Samtools (v1.3) (72) and unix awk commands. PCR duplicates were then removed using PicardTools (v1.83) and only uniquely mapped reads were retained. Insert sizes were obtained from the respective BAM files using PicardTools (v1.83). All samples displayed the expected periodicity of DNA winding around nucleosomes in genome DNA regions. A custom Perl script was used to generate smoothened genome browser tracks in BigWig format for data visualisation on the UCSC Genome Browser.

### Transcription factor binding analysis

Avi-tagged Sox10 biotin ChIP sequencing and Cut and Run samples were demultiplexed and the resulting files were merged. Read quality evaluation was performed using FastQC v0.11.4 (70). Adaptor sequences were trimmed using TrimGalore v0.4.1 and then mapped to the chicken genome galGal5 assembly using bowtie2 v2.3.5 (73). Only aligned pairs were retained based on binary alignment map (BAM) flags, and mitochondrial reads were removed from the BAM file using Samtools (v1.3) (72). PCR duplicates were then removed using PicardTools (v1.83) and only uniquely mapped reads were retained. A custom Perl script was used to generate smoothened genome browser tracks in BigWig format for data visualisation on the UCSC Genome Browser. Differential binding analysis was carried out with the DiffBind R package (v1.10.2).

### Data availability

ChIP-seq and ATAC-seq datasets associated to *in vitro* derived human cranial neural crest cells (44) have been retrieved from the NCBI GEO repository under accession ID GEO: GSE70751. ATAC-seq and ChIP-seq from avian neural crest cells (34, 41) were retrieved from GEO IDs GSE121527 and GSE125711, respectively. The raw and processed data generated in this study have been submitted to GEO under accession ID GSEXXXXX.

## Supplementary Figures

**Fig. S1.**
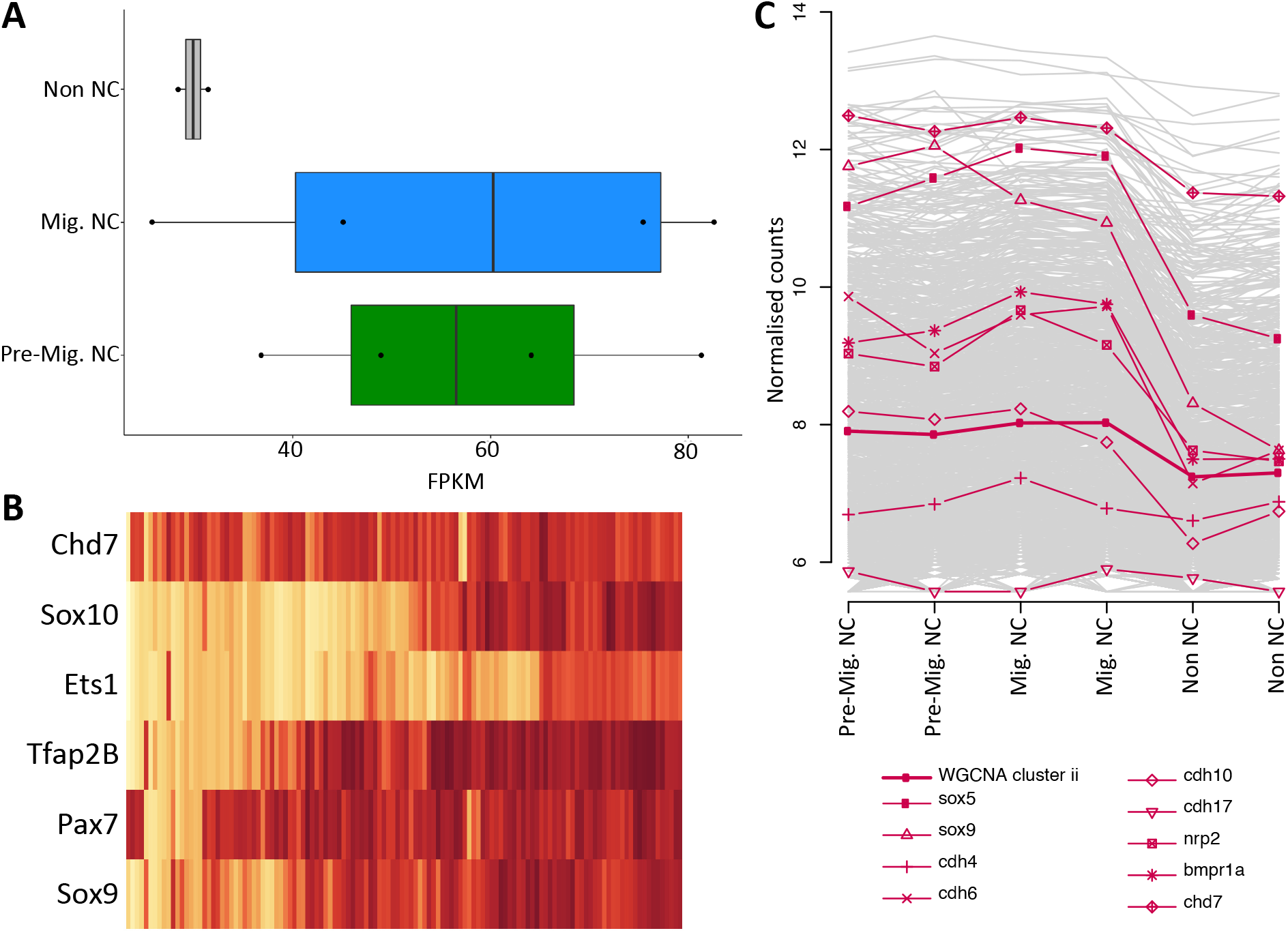
*Chd7* enrichment in neural crest RNA-seq data. (A) Differential expression analysis of RNA-seq data sets obtained from cranial neural crest cells and surrounding non-neural crest cells showed *Chd7* is elevated in the neural crest at pre-migratory (HH9) and early migratory (HH10) stages (p-adj 1.512-01 and 9.46-01 respectively). (B) Co-expression of *Chd7* and other selected genes in neural crest single-cell (Smartseq2) RNA-seq data. (C) WGCNA plot showing clustering of *Chd7* with other neural crest regulators. Data and images taken from [32].

**Fig. S2.**
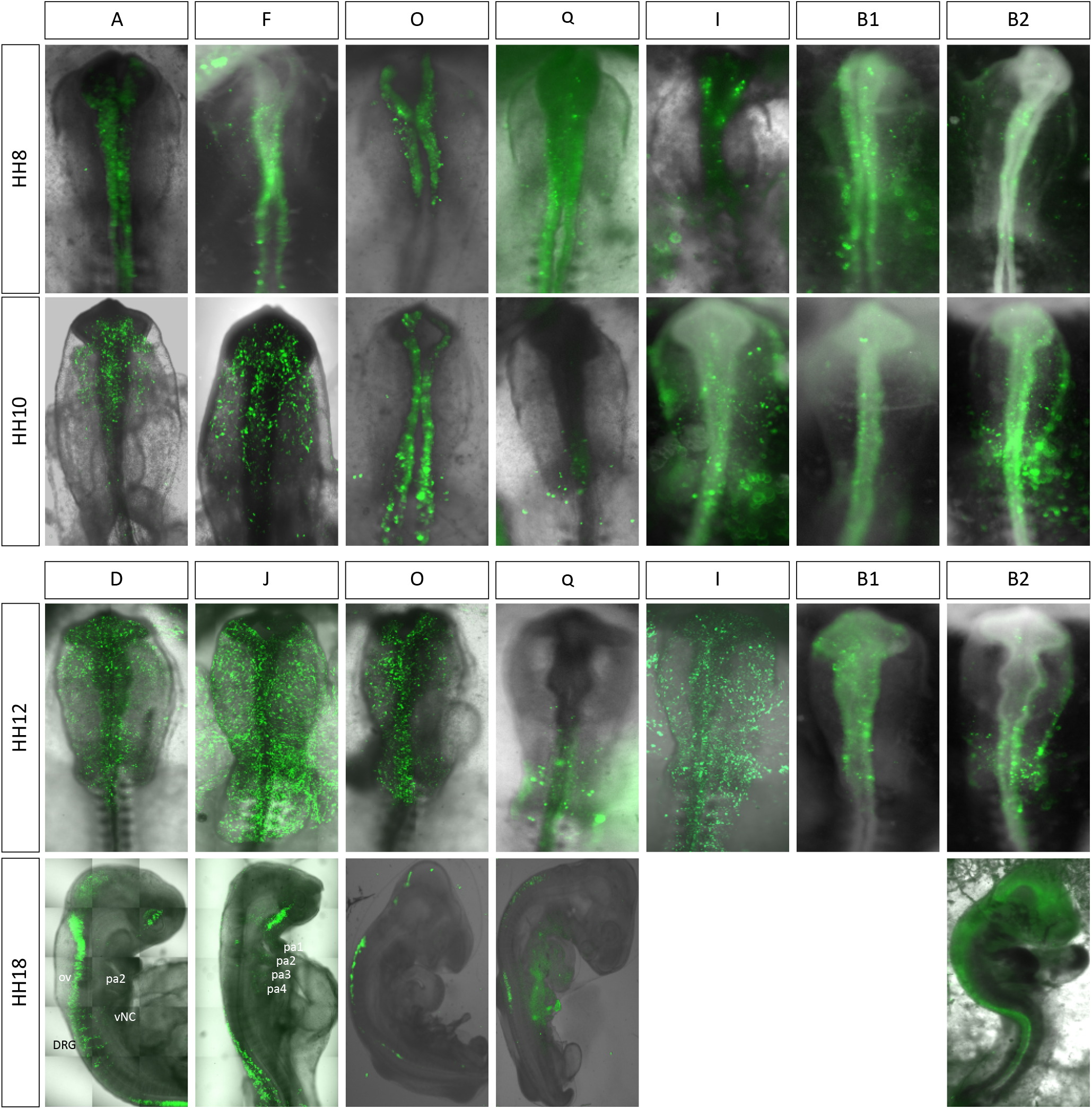
*In vivo* activity of *Chd7* enhancers. Chick *Chd7* enhancers, as indicated by top boxes, shown at alternative stages. Additional enhancers; B1, B2, I, J, O, Q are also shown.

**Fig. S3.**
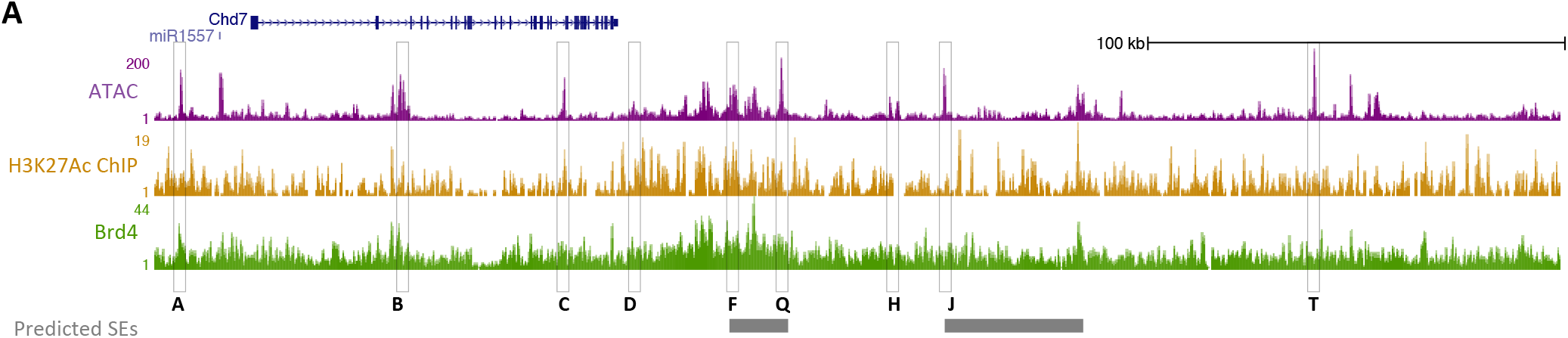
*Chd7* super-enhancer. (A) Genome browser view of ATAC (purple) and ChIP, H3K27Ac (yellow) and Brd4 (green), data from chick neural crest cells. Superenhancer regions, predicted by the ROSE algorithm, are shown in grey. Some *Chd7* enhancers are also shown for context.

**Fig. S4.**
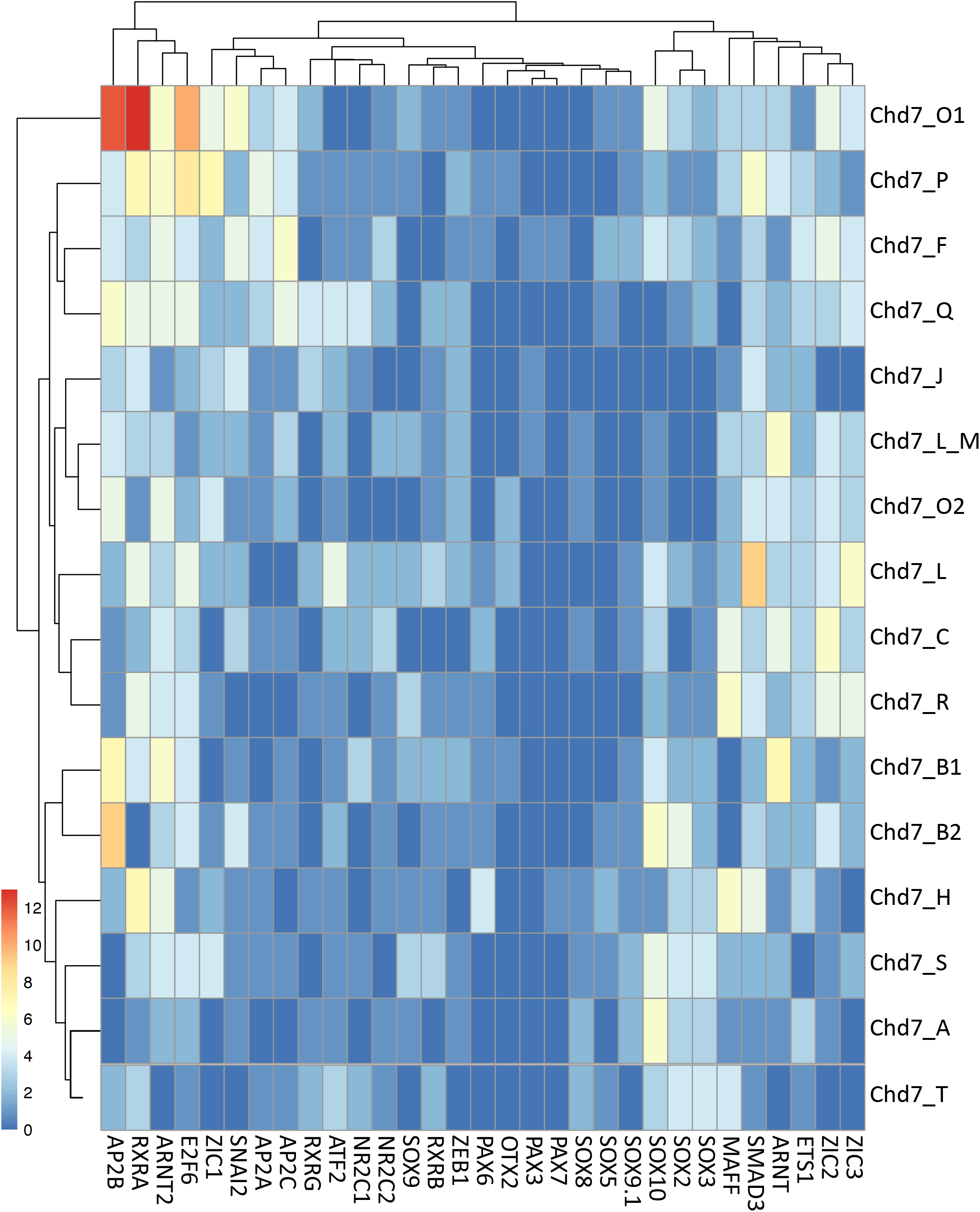
Transcription factor motif sites within *Chd7* enhancers. Heatmap of predicted transcription factor motifs identified in *Chd7* enhancers using HOMER.

